# Epigenome-wide association in adipose tissue from the METSIM cohort

**DOI:** 10.1101/223495

**Authors:** Luz D. Orozco, Colin Farrell, Christopher Hale, Liudmilla Rbi, Arturo Rinaldi, Mete Civelek, Calvin Pan, Larry Lam, Dennis Montoya, Chantle Edillor, Marcus Seldin, Karen L Mohlke, Steve Jacobsen, Johanna Kuusisto, Markku Laakso, Aldons J Lusis, Matteo Pellegrinil

**Affiliations:** Department of Molecular, Cell and Developmental Biology, University of California Los Angeles, Los Angeles, CA, 90095, USA; Departments of Human Genetics, Medicine, and Microbiology, David Geffen School of Medicine, University of California Los Angeles, Los Angeles, CA, 90095, USA; Department of Genetics, University of North Carolina, Chapel Hill, NC 27599, USA; Department of Biostatistics and Center for Statistical Genetics, University of Michigan, Ann Arbor, Michigan 48109, USA; Eli & Edythe Broad Center of Regenerative Medicine & Stem Cell Research, University of California Los Angeles, Los Angeles, California 90095, USA; Howard Hughes Medical Institute, University of California Los Angeles, Los Angeles, California 90095, USA; Institute of Clinical Medicine, Internal Medicine, University of Eastern Finland and Kuopio University Hospital

## Abstract

Most epigenome-wide association studies to date have been conducted in blood. However, metabolic syndrome is mediated by a dysregulation of adiposity and therefore it is critical to study adipose tissue in order to understand the effects of this syndrome on epigenomes. To determine if natural variation in DNA methylation was associated with metabolic syndrome traits, we profiled global methylation levels in subcutaneous abdominal adipose tissue. We measured association between 32 clinical traits related to diabetes and obesity in 201 people from the Metabolic Syndrome In Men cohort. We performed epigenome-wide association studies between DNA methylation levels and traits, and identified associations for 13 clinical traits in 21 loci. We prioritized candidate genes in these loci using eQTL, and identified 18 high confidence candidate genes, including known and novel genes associated with diabetes and obesity traits. Using methylation deconvolution, we examined which cell types may be mediating the associations, and concluded that most of the loci we identified were specific to adipocytes. We determined whether the abundance of cell types varies with metabolic traits, and found that macrophages increased in abundance with the severity of metabolic syndrome traits. Finally, we developed a DNA methylation based biomarker to assess type II diabetes risk in adipose tissue. In conclusion, our results demonstrate that profiling DNA methylation in adipose tissue is a powerful tool for understanding the molecular effects of metabolic syndrome on adipose tissue, and can be used in conjunction with traditional genetic analyses to further characterize this disorder.

## INTRODUCTION

Metabolic syndrome traits such as obesity, dyslipidemia, insulin resistance, and hypertension underlie the common forms of atherosclerosis, type 2 diabetes (T2D) and heart failure, which together account for the majority of deaths in Western populations. Metabolic syndrome affects 44% of adults over the age of 50 in the US, and people affected with metabolic syndrome have higher risk of heart attacks, diabetes and stroke (1). Numerous studies have investigated the genetic basis of metabolic syndrome traits such as diabetes (2), and accumulating evidence suggests that epigenetics is associated with these phenotypes (3, 4).

Methylation of DNA cytosine bases is evolutionarily conserved and plays important roles in development, cell differentiation, imprinting, X-chromosome inactivation, and regulation of gene expression. Aberrant DNA methylation in mammals is associated with both rare and complex traits including cancer, aging (5), and imprinting disorders such as Prader-Willi syndrome. Recent studies have demonstrated that much like genome sequence variation, DNA methylation is variable among individuals in human (6), plant (7) and mouse (8) populations. Moreover, differences in DNA methylation of cytosines are in part heritable and controlled by genetics both in *cis* and in *trans.* However, sex and environmental factors such as smoking and diet can also influence DNA methylation differences, leading to changes in methylation levels over an individual’s lifetime (9).

DNA methylation states have been shown to be associated with biological processes underlying metabolic syndrome, including obesity, hypertension and diabetes (10). Environment-induced changes in DNA methylation have also been associated with fetal origins of adult disease (11), and alterations in maternal diet during pregnancy can affect the methylation levels of the placenta, inducing transcriptional changes in key metabolic regulatory genes (12, 13). Recent studies have also shown that diet-induced obesity in adults affects methylation of obesogenic genes such as leptin (14), *Scd1* (15), and *LPK* (16).

Similar to genome-wide association studies (GWAS), *epi*genome-wide association studies (EWAS) aim to identify candidate genes for traits by using epigenetic factors instead of SNP genotypes in the association model. EWAS have recently identified associations for gene expression and protein levels in humans (6), and complex traits such as bone mineral density, obesity, and insulin resistance in mice (17). However, to date, most EWAS studies have been carried out in blood, which is the tissue that is most readily collected for large scale studies in humans.

By contrast, in this study we examined the association of DNA methylation with metabolic traits in humans using adipose tissue samples from the Metabolic Syndrome in Men (METSIM) cohort. Metabolic syndrome is characterized by a clustering of three or more of the following conditions: elevated blood pressure, elevated serum triglycerides, elevated blood sugar, low HDL levels, and abdominal obesity. As adipose is known to be a central organ in metabolic syndrome manifestation, adipose tissue should be one of the most relevant for defining and studying metabolic syndrome traits (18) The METSIM cohort has been thoroughly characterized for longitudinal clinical data of metabolic traits including a 3-point oral glucose tolerance test, cardiovascular disorders, diabetes complications, drug and diet questionnaire, as well as high density genotyping, and genome-wide expression in adipose (19, 20). We performed EWAS on clinical traits using reduced representation bisulfite sequencing data and identified 51 significant associations for metabolic syndrome traits, corresponding to 21 loci. These associations include previously known genes, *FASN* (21–23) and *RXRA* (24–26), as well as loci harboring 22 new candidate genes for diabetes and obesity in humans. We identify the types of cells that are likely to be mediating these associations, and conclude that adipocytes are involved. We also examine the abundance of cell types and show that macrophages increase with the severity of metabolic syndrome traits. Finally, we develop a biomarker for to assess type II diabetes status in adipose tissue. Our results demonstrate that DNA methylation profiling is both useful and complementary to GWAS for characterizing the molecular and cellular basis of metabolic syndrome.

## RESULTS

### METSIM COHORT

The METSIM cohort consists of 10,197 men from Kuopio Finland between 45-73 years of age. Laakso and colleagues (20) have characterized this cohort for numerous clinical traits involved in diabetes and obesity, and genome-wide expression levels in adipose tissue biopsies (19). In this study, we examined 32 clinical traits related to metabolic syndrome (Table S1), adipose tissue expression levels using microarrays, and DNA methylation profiles from adipose tissue biopsies in 201 individuals from the METSIM cohort.

### DNA METHYLATION OF ADIPOSE BIOPSIES

To examine methylation patterns in the METSIM cohort we constructed reduced representation bisulfite sequencing libraries (RRBS) from adipose tissue biopsies, corresponding to 228 individuals. The sequences obtained from RRBS libraries are enriched in genes and CpG islands, and cover 4.6 million CpGs out of the ∼ 30 million CpGs in the human genome (∼ 15%). We sequenced the libraries using the Illumina HiSeq platform and obtained on average 34.3 + /-6.7 million reads per sample. We aligned the data to the human genome using BSMAP (27) and obtained on average 21.9 + /-4.6 million uniquely aligned reads per sample (Figure S1A), corresponding to 64% average mappability (Figure S1B), and 19x average coverage (Figure S1B). We focused our analyses on CpGs, since CHG and CHH (H = A, C or T) methylation in mammals is on average only 1-2%, which makes it difficult to detect significant variation in our samples (17, 28). We and others have previously validated RRBS data relative to traditional bisulfite sequencing by cloning DNA fragments into bacterial colonies followed by Sanger sequencing and found a high degree of concordance between RRBS and traditional bisulfite sequencing results in mice (8, 12) and humans (29). RRBS shows limited overlap with the Illumina 450k arrays, a small study (n = 11) found an overlap between 24,000-120,000 CpG sites (30).

We filtered our dataset for CpGs with at least 10x coverage, and present in at least 75% of the samples, corresponding to 2,320,297 CpGs. However, the methylation state of individual CpGs may be subject to stochastic variation or measurement error, and we observed a single or a few outlier samples with methylation levels that are very different from the rest of the population (Figure S1C). This variability is likely to lead to spurious associations between methylation and traits, and we observed that was indeed the case when we performed EWAS using individual CpGs. In contrast to individual CpGs, the methylation level of a CpG methylation region (a unit comprised of several CpGs) is a much more robust measure of DNA methylation levels (for example see Figure S1D). Methylation regions are likely a more biologically relevant genomic unit than individual CpGs, and methylation levels of proximal CpGs tend to be correlated over distances of a few hundred bases to 1kb, roughly the typical size of CpG islands (31). Therefore, we defined 149,191 methylation regions, where each region is defined as the average methylation of multiple CpGs that are near each other and highly correlated. We require that a region have a minimum of 2 CpGs whose methylation is correlated (Pearson’s r > = 0.9), and the region has a maximum size of 3kb. The distribution of methylation levels for these regions is shown in Figure S1E, with average methylation levels of 58.4% + /-4.7. The range of methylation across all individuals can vary between 0 and 100%. The average range was 33% for individual CpGs, and 25% for methylation regions (Figure S1F). These regions are located throughout the genome with no major gaps in coverage, with the exception of centromeric and acrocentric regions, and in the y-chromosome where there was minimal coverage (Figure S2).

### EWAS

We performed epigenome-wide association (EWAS) between CpG methylation regions and 32 clinical traits related to obesity and diabetes, including body weight, body mass index (BMI), body fat percentage, oral glucose tolerance test (OGTT), glucose and insulin measurements (Table S1). All clinical traits were transformed using inverse normal transformation (see Methods), as is common practice for genome-wide association studies of quantitative traits (32). We used the linear mixed model package pyLMM to determine associations between DNA methylation patterns and phenotypes. Others and we have previously demonstrated that this approach corrects for spurious associations due to population structure (17, 33) and tissue heterogeneity (34). Associations were considered significant if the *p*-value for the association was below 1×10^-7^, based on the Bonferroni correction for the number of CpG regions tested.

In total, we found 51 significant associations, corresponding to 21 distinct methylation loci and 15 unique phenotypes (Figure 1 and Table 1) where the *p*-value was below 1×10^-7^. Of the 21 distinct loci, 15 methylation loci were intragenic, and 6 loci were intergenic. The distance between intergenic loci and nearby flanking genes ranged between 23kb and 440kb. Candidate genes listed for each association in Table 1 correspond to the gene itself for intragenic associations, and the two nearest flanking genes by distance for intergenic associations, with the distance between the locus and each flanking gene listed for intergenic associations (Table 1). Figure 1 summarizes the genomic distribution of all EWAS hits, where each dot represents an association between a phenotype and a methylation region.

**Table 1:** Clinical trait EWAS

| Table 1: Clinical trait EWAS |  |  |  |  |  |  |  |  |  |  |  |
| --- | --- | --- | --- | --- | --- | --- | --- | --- | --- | --- | --- |
| EWAS locus | Chr | Bp start | Bp end | Candidate Genes(s) | cis-eQTL | Intra/inter-genic | EWAS clinical trait | EWAS Pval | EWAS beta (effect size) | EWAS Inflation factor $\lambda$ | Methylation range |
| Met1 | 2 | 34545726 | 34545903 | *LINC01317, LINC01320* |  | Intergenic, 23kb, 356kb | MATSUDA insulin sensitivity index | 7.53E-19 | 39.30 | 0.99 | 16.10% |
| Met2 | 2 | 47979865 | 47980000 | *MSH2, MSH6* | MSH2 cis-eQTL (p=3.3e-94, rs2303425 ), and MSH6 cis-eQTL (p=1.1e-13, rs2134056) | Intergenic, 350kb, 30kb | Waist circumference | 3.26E-08 | -4.74 | 0.99 | 68.52% |
| Met2 |  |  |  |  |  |  | Fat mass (%) | 6.08E-10 | -5.27 | 1.00 |  |
| Met2 |  |  |  |  |  |  | Fat free mass (%) | 6.11E-10 | 5.27 | 0.99 |  |
| Met2 |  |  |  |  |  |  | Body mass index | 7.75E-09 | -5.04 | 1.01 |  |
| Met3 | 2 | 65064793 | 65064867 | *AFTPH, SLC1A4* | SLC1A4 cis-eQTL (p=1.6e-10, chr2:65252385) | Intergenic, 244kb, 150kb | Waist circumference | 4.92E-08 | -4.44 | 0.99 | 58.28% |
| Met3 |  |  |  |  |  |  | Fat mass (%) | 3.89E-09 | -4.82 | 1.00 |  |
| Met3 |  |  |  |  |  |  | Fat free mass (%) | 3.57E-09 | 4.83 | 0.99 |  |
| Met3 |  |  |  |  |  |  | OGTT fasting plasma insulin | 2.25E-09 | -4.85 | 0.97 |  |
| Met3 |  |  |  |  |  |  | Body mass index | 5.71E-10 | -5.05 | 1.01 |  |
| Met3 |  |  |  |  |  |  | MATSUDA insulin sensitivity index | 8.30E-09 | 4.76 | 0.99 |  |
| Met3 |  |  |  |  |  |  | HOMAIR Insulin resistance index based on HOMA | 2.19E-09 | -4.85 | 0.97 |  |
| Met4 | 4 | 7774215 | 7774379 | *AFAP1* | AFAP1 cis-eQTL (p=1.16e-16, rs34072960) | Intragenic | Fat mass (%) | 4.43E-08 | 6.59 | 1.00 | 78.69% |
| Met4 |  |  |  |  |  |  | Fat free mass (%) | 4.42E-08 | -6.59 | 0.99 |  |
| Met4 |  |  |  |  |  |  | Body mass index | 5.57E-08 | 6.65 | 1.01 |  |
| Met5 | 5 | 173132627 | 173132883 | *BOD1, CPEB4* | BOD cis-eQTL (p=1.3e-6, rs60748211), CPEB4 cis-eQTL (p=2.4e-174, rs72812818) | Intergenic, 88kb, 182kb | OGTT fasting plasma insulin | 1.97E-08 | 5.95 | 0.97 | 77.70% |
| Met5 |  |  |  |  |  |  | OGTT 120 min plasma insulin | 2.45E-09 | 6.07 | 0.98 |  |
| Met5 |  |  |  |  |  |  | MATSUDA insulin sensitivity index | 4.42E-10 | -6.81 | 0.99 |  |
| Met5 |  |  |  |  |  |  | HOMAIR Insulin resistance index based on HOMA | 1.04E-08 | 6.10 | 0.97 |  |
| Met6 | 6 | 3132000 | 3132119 | *BPHL* | BPHL cis-eQTL (p=5.0e-81, rs7765391) | Intragenic | Weight | 9.22E-08 | 11.16 | 0.99 | 19.17% |
| Met7 | 9 | 137263485 | 137263573 | *RXRA* | RXRA cis-eQTL (p=1.0e-09, rs62576325) | Intragenic | Waist circumference | 6.08E-08 | 3.41 | 0.99 | 85% |
| Met8 | 10 | 75600604 | 75600822 | *CAMK2G* | CAMK2G cis-eQTL (p=3.9e-28, rs2675671) | Intragenic | OGTT 120 min plasma proinsulin | 9.54E-08 | 18.43 | 0.96 | 10.40% |
| Met9 | 10 | 100183346 | 100183441 | *HPS1* | HPS1 cis-eQTL (p=1.1e-53, rs701801) | Intragenic | Waist circumference | 1.55E-11 | 10.02 | 0.99 | 30% |
| Met10 | 11 | 65248602 | 65248737 | *FRMD8, SCYL1* |  | Intergenic, 67kb, 43kb | OGTT fasting plasma insulin | 6.84E-08 | -16.14 | 0.97 | 30% |
| Met10 |  |  |  |  |  |  | HOMAIR Insulin resistance index based on HOMA | 5.45E-08 | -16.25 | 0.97 |  |
| Met11 | 12 | 42883183 | 42883320 | *PRICKLE1* |  | Intragenic | OGTT 30 min plasma insulin | 1.70E-19 | -17.20 | 0.81 | 17.64% |
| Met12 | 12 | 113734359 | 113734579 | *TPCN1* | TPCN1 cis-eQTL (p=1.4e-6, rs2004720) | Intragenic | Weight | 7.24E-08 | 5.72 | 0.99 | 13.24% |
|  |  |  |  |  |  |  | Glomerular filtration rate (eGFRcockroft) | 6.24E-08 | 5.61 | 0.97 | 71.39% |
| Met13 | 17 | 2893657 | 2893829 | *RAP1GAP2* |  | Intragenic | OGTT fasting plasma insulin | 2.93E-08 | -11.03 | 0.97 | 30.25% |
| Met13 |  |  |  |  |  |  | HOMAIR Insulin resistance index based on HOMA | 1.96E-08 | -11.16 | 0.97 |  |
| Met14 | 17 | 40913548 | 40913600 | *RAMP2* |  | Intragenic | OGTT insulin area under the curve | 6.51E-29 | 12.06 | 0.99 | 32.53% |
| Met15 | 17 | 80041514 | 80042067 | *FASN* | FASN cis-eQTL (p=9.3e-10, rs4239015 ) | Intragenic | Weight | 4.07E-09 | 19.42 | 0.99 | 12.18% |
| Met15 |  |  |  |  |  |  | Waist circumference | 8.54E-08 | 17.31 | 0.99 |  |
| Met15 |  |  |  |  |  |  | Fat mass (%) | 1.36E-08 | 18.29 | 1.00 |  |
| Met15 |  |  |  |  |  |  | Fat free mass (%) | 1.40E-08 | -18.27 | 0.99 |  |
| Met15 |  |  |  |  |  |  | Body mass index | 3.05E-10 | 21.57 | 1.01 |  |
| Met16 | 17 | 80046324 | 80047195 | *FASN* |  | Intragenic | Body mass index | 1.18E-08 | 16.61 | 1.01 | 20.05% |
| Met17 | 17 | 80050198 | 80050365 | *FASN* |  | Intragenic | Body mass index | 2.48E-10 | 8.01 | 1.01 | 50.31% |
| Met18 | 17 | 80051500 | 80053080 | *FASN* |  | Intragenic | Weight | 4.08E-09 | 7.95 | 0.99 | 67.62% |
| Met18 |  |  |  |  |  |  | Waist circumference | 4.29E-10 | 8.33 | 0.99 |  |
| Met18 |  |  |  |  |  |  | Fat mass (%) | 1.03E-08 | 7.85 | 1.00 |  |
| Met18 |  |  |  |  |  |  | Fat free mass (%) | 9.81E-09 | -7.86 | 0.99 |  |
| Met18 |  |  |  |  |  |  | OGTT fasting plasma insulin | 1.08E-08 | 7.66 | 0.97 |  |
| Met18 |  |  |  |  |  |  | OGTT 120 min plasma insulin | 5.42E-09 | 8.03 | 0.98 |  |
| Met18 |  |  |  |  |  |  | Body mass index | 1.37E-11 | 9.44 | 1.01 |  |
| Met18 |  |  |  |  |  |  | MATSUDA insulin sensitivity index | 1.79E-08 | -8.05 | 0.99 |  |
| Met18 |  |  |  |  |  |  | HOMAIR Insulin resistance index based on HOMA | 3.57E-09 | 7.94 | 0.97 |  |
| Met19 | 19 | 1148096 | 1148232 | *SBNO2* |  | Intragenic | Weight | 3.20E-08 | 8.54 | 0.99 | 40.68% |
| Met19 |  |  |  |  |  |  | Body mass index | 7.58E-08 | 8.49 | 1.01 |  |
| Met20 | 21 | 16661536 | 16661714 | *NRIP1, USP25* | NRIP1 cis-eQTL (p=6.6e-85, rs2178895) | Intergenic, 224kb, 440kb | Body mass index | 3.27E-08 | 8.14 | 1.01 | 86.15% |
| Met21 | 22 | 28151964 | 28152032 | *MN1* |  | Intragenic | OGTT 120 min plasma insulin | 9.16E-08 | 7.64 | 0.98 | 30.95% |

**Figure 1:**
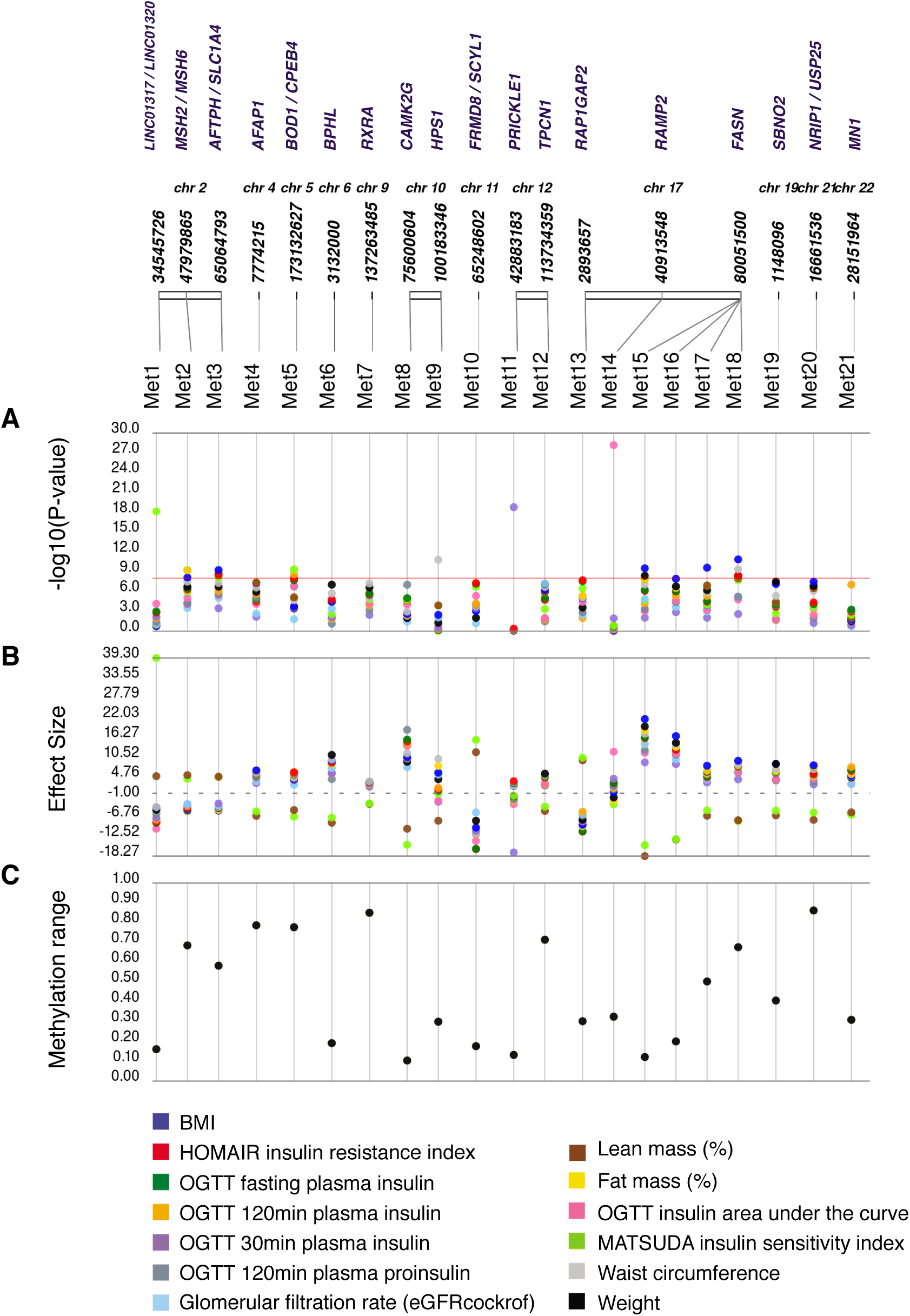
Epigenome-wide association of metabolic clinical traits. Association between DNA CpG methylation and clinical traits. (A) “PheWAS” plot showing association of each of the methylation loci (Met 1-21) and the clinical traits. The genomic location of the CpG is on the x-axis and the association significance is on the y-axis. Different colors represent different traits. (B) The effect size for each association shown in (A). (C) For each methylation locus on the x-axis, the range of methylation across individuals in the population is shown on the y-axis.

Some may argue that a significance threshold of 1×10^-7^ is insufficiently low, since we tested for 32 traits. Of the 51 associations described above, 26 associations would remain significant using a Bonferroni correction (*p*<1×10^-8^) which accounts for both CpG regions and the 32 traits. However, we believe the additional Bonferroni correction for 32 traits would be too stringent given that several of the traits are not independent, for example BMI and fat mass, or plasma insulin and glucose levels. Alternatively, 40 associations would remain significant at the commonly used GWAS significance threshold (*p*<5×10^-8^).

We found no evidence of inflation in our EWAS results, where the inflation factor lambda was on average 0.99, and maximum of 1.01 (Table 1). Sample EWAS *p*-value distributions and ggplots are shown in Figure S3A-C.

### CANDIDATE GENES

We initially identified a total of 24 candidate genes and non-coding RNAs by proximity to an EWAS signal (Table 1). To prioritize candidate genes, we examined adipose expression associations from 770 individuals of the METSIM cohort previously published by our laboratories (19, 35). We asked if there were significant expression quantitative trait loci that overlapped with the methylation loci identified in the EWAS. We narrowed down the candidate gene list from 24 to 18 (75%) high confidence candidate genes that had significant cis-eQTL in adipose tissue samples from the METSIM cohort (Table 1). The *cis -* eQTL were significant for the candidate gene reported.

We identified 3 loci where multiple clinical traits mapped to the same methylation region. These loci include chromosome 17 at the *FASN* gene (Figure 2A-B), in chromosome 2 near *SLC1A4*, and in chromosome 5 near *CPEB4* (Figure 1, Table 1). The *FASN* gene has a cis-eQTL (*p*=9.3×10^-10^, Figure 2C), suggesting that genetic variation in the population affects expression levels of this gene. One of the traits associated with this locus is BMI, and we observed a positive correlation between methylation levels in the *FASN* locus and BMI (Figure 2D). Remarkably, the observed correlation of 0.4 suggests that the methylation of FASN alone is able to capture 16% of the variation in BMI observed in our cohort. Moreover, we also observed an inverse correlation between methylation and *FASN* expression in adipose tissue biopsies (Figure 2E), and an inverse correlation between *FASN* expression and BMI (Figure 2F).

**Figure 2:**
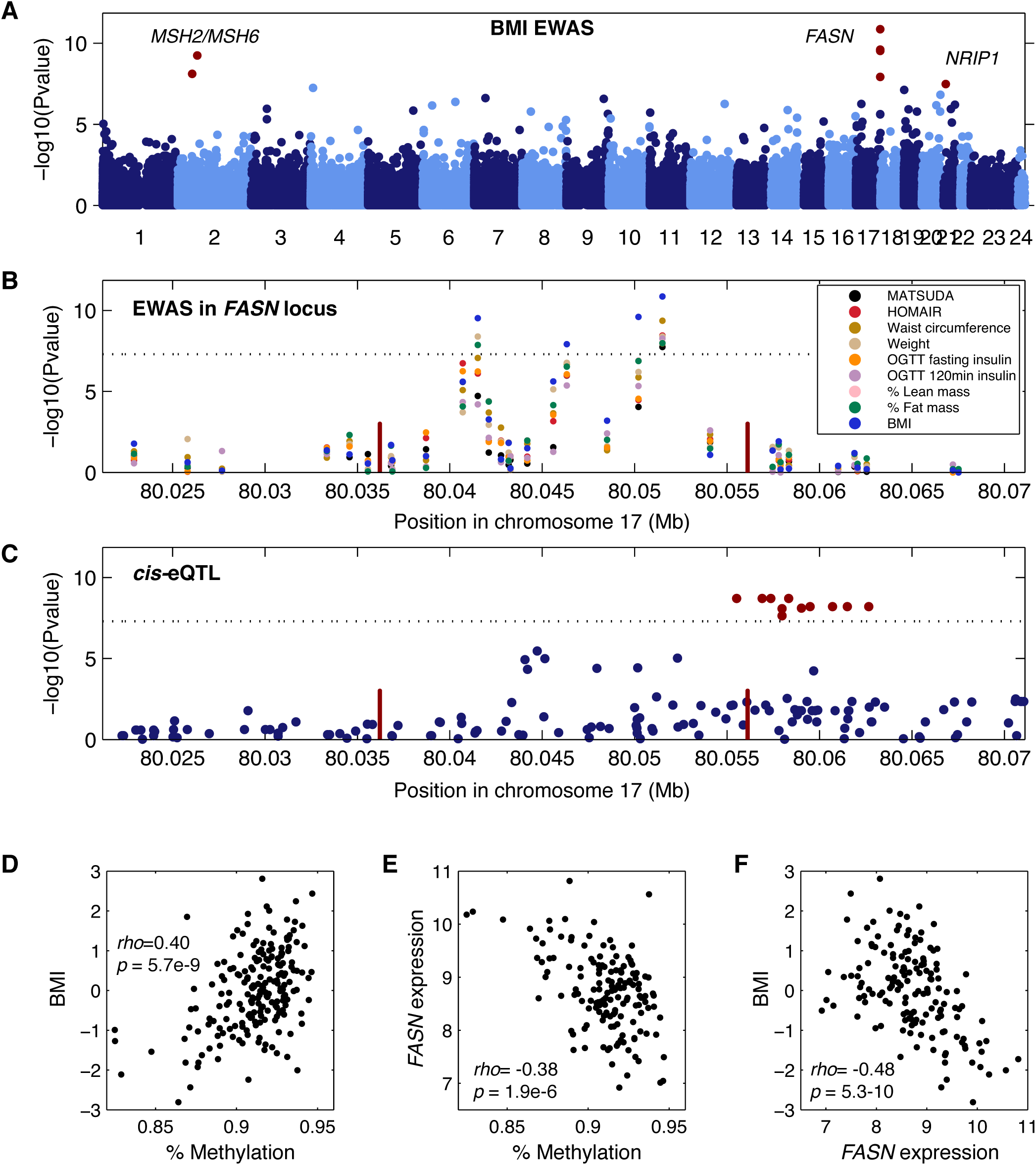
FASN is associated with multiple clinical traits. (A) Manhattan plot showing EWAS results for BMI. Each dot represents a CpG region with the genomic location of each CpG region on the x-axis and chromosomes shown in alternating colors. The association significance is on the y-axis and significant hits are shown as red dots. (B) Association results for multiple phenotypes near the FASN locus. Each dot represents a different association to a CpG region and different colored points represent distinct clinical traits. The genomic location of each CpG region is on the x-axis and the association significance is on the y-axis. Red vertical bars denote the transcription start and end of FASN. The dotted significance threshold line is drawn at 5×10-8. (C) cis-eQTL results for FASN expression in adipose tissue biopsies. Each dot represents a SNP. The genomic location of each SNP is on the x-axis and the association significance is on the y-axis. Significant SNPs are shown as red dots. Red vertical bars denote the transcription start and end of FASN. The dotted significance threshold line is drawn at 5×10-8. (D)-(F) Each point represents an individual in the cohort, showing correlation between (D) methylation levels for the peak associated CpG region and BMI, (E) methylation levels for the peak CpG region and expression of FASN, and (F) expression of FASN and BMI.

A second locus is located upstream of *SLC1A4* and was associated with waist circumference, lean mass, fat mass, plasma insulin levels, BMI, and indices of insulin resistance and insulin sensitivity MATSUDA, and HOMAIR (Figure 3A-B). *SLC1A4* has a cis-eQTL (*p*=1.6×10^-10^, Figure 3C), and we observed an inverse correlation between methylation at this locus and the insulin resistance index HOMAIR (Figure 3D), an inverse correlation between methylation and expression of *SLC1A4* (Figure 3E), and a positive correlation between *SLC1A4* expression and HOMAIR (Figure 3F). A third locus is located upstream of *CPEB4*, and was associated with basal plasma insulin levels, OGTT plasma insulin, and the indices of insulin resistance and insulin sensitivity MATSUDA, and HOMAIR (Table 1). A *cis -* eQTL for *CPEB4* expression in adipose tissue biopsies (p> = 2.4×10^-174^) makes this gene strong candidate gene for this locus.

**Figure 3:**
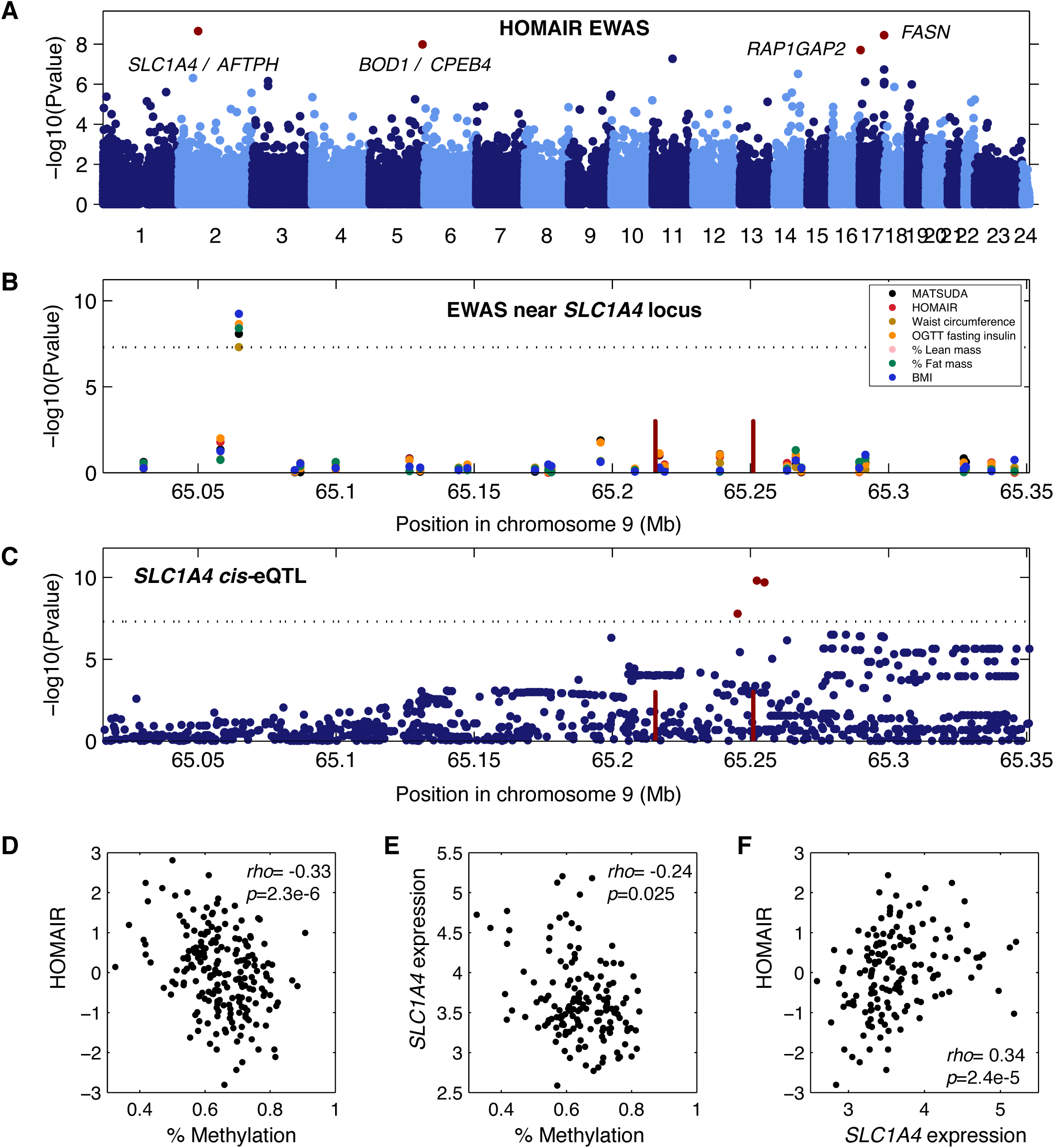
SLC1A4 is associated with multiple clinical traits. (A) Manhattan plot showing EWAS results for insulin resistance index HOMAIR. Each dot is a CpG region, the genomic location of each CpG region is on the x-axis with chromosomes shown in alternating colors, the association significance is on the y-axis, significant hits are shown as red dots. (B) Association results for multiple phenotypes near the SLC1A4 locus. Each dot represents a different association to a CpG region, different colored points represent distinct clinical traits, the genomic location of each CpG region is on the x-axis, the association significance is on the y-axis. Red vertical bars denote the transcription start and end of SLC1A4. The dotted significance threshold line is drawn at 5×10^-8^. (C) cis-eQTL results for SLC1A4 expression in adipose tissue biopsies. Each dot represents a SNP, the genomic location of each SNP is on the x-axis, the association significance is on the y-axis, significant SNPs are shown as red dots. Red vertical bars denote the transcription start and end of SLC1A4. The dotted significance threshold line is drawn at 5×10^-8^. (D)-(F) Each point represents an individual in the cohort, showing correlation between (D) methylation levels for the peak associated CpG region and HOMAIR, (E) methylation levels for the peak CpG region and expression of SLC1A4, and (F) expression of SLC1A4 and HOMAIR.

### CELL TYPE DECOMPOSITION OF ADIPOSE TISSUE

Whenever we examine molecular phenotypes such as DNA methylation and gene expression in tissues, the question arises, what cell-types within the tissue are responsible for the signal we observe? We know that subcutaneous adipose tissue is composed primarily of adipocytes, but also contains endothelial cells and immune cells such as resident and infiltrating macrophages. Moreover, we know that obese individuals show increased macrophage content in their adipose tissue, and hence that heterogeneity in people’s phenotypes can influence cell-type composition in the adipose tissue biopsies (36). To examine macrophage content in the adipose tissue biopsies, we examined expression levels of genes expressed in adipocytes, namely *PPARG, CFD, ADIPOQ, FABP4, CIDEC, LEP*, and *TNMD*, and genes highly expressed in macrophages including *TLR1, TLR2, TLR3, TLR4, ABCG, IL10*, and *TNF.* We found high expression levels of adipocyte-specific genes and low expression of macrophage-specific genes (Figure S3D). These results suggest that there is minimal macrophage content in the adipose biopsies. However, the genes selected may not fully reflect the transcriptome of adipocytes and macrophages, or additional cell-types that may be present in adipose tissue.

To further explore the contribution of different cell-types to the METSIM adipose tissue biopsies, we performed cell-type deconvolution using BS-seq methylation data from our samples and from multiple reference cell-types including adipocytes, endothelial cells, macrophages, neutrophils, NK -, T -, and B-cells. Using this approach, we can determine the relative content of different cell types by comparing DNA methylation at cell-specific methylation markers in our test samples, to DNA methylation signatures derived from purified cell types (see Methods). Consistent with our previous analysis of gene expression in adipose-and macrophage-specific genes, we found that the highest cell-type represented in our adipose biopsies was indeed adipocyte (Figure 4A), but we also found evidence of macrophage and neutrophil content.

**Figure 4:**
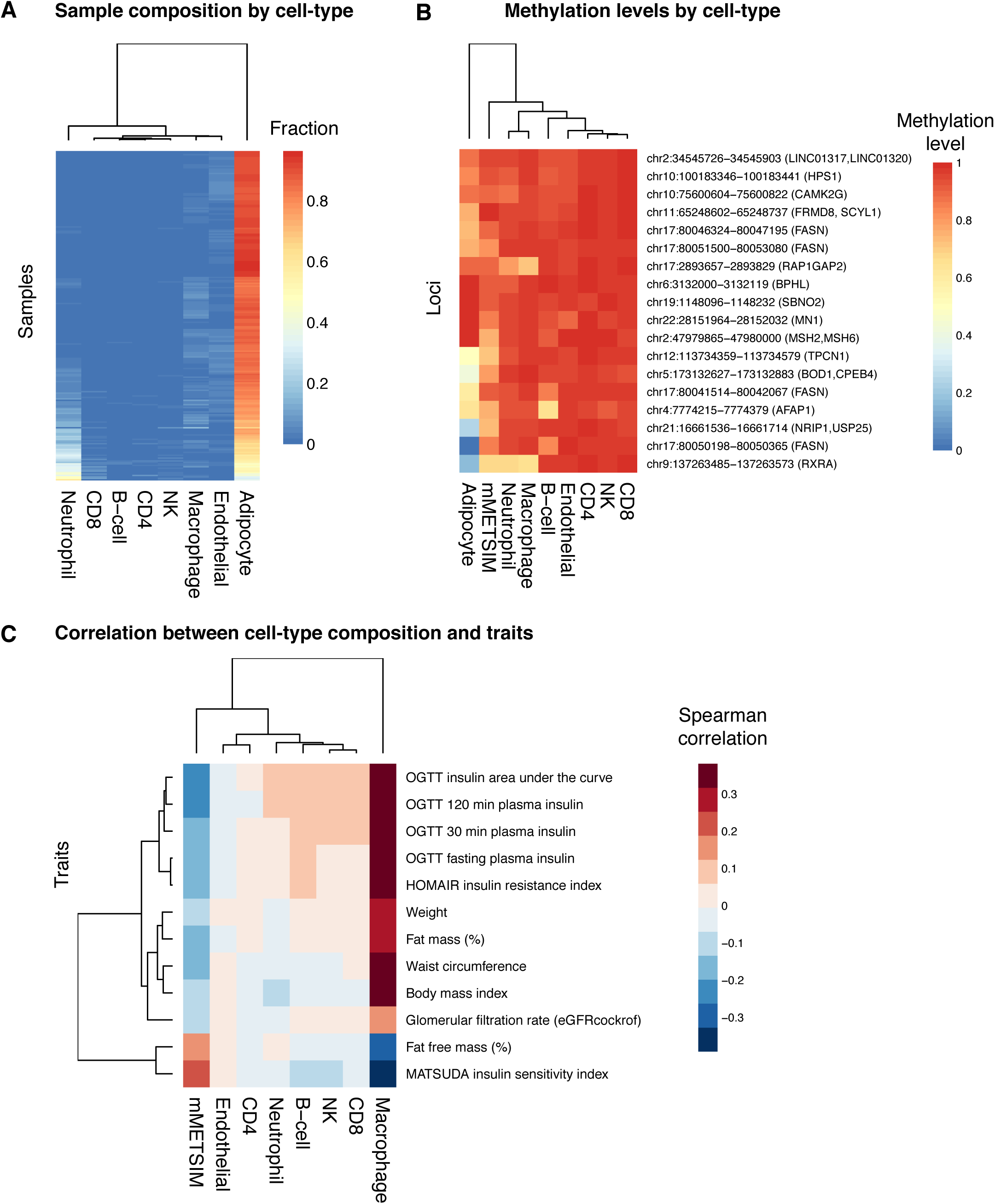
Cell-type deconvolution. (A) Sample composition by cell-type is shown for different cell types (columns), across all METSIM samples (rows). The color in the heatmap represents the relative fraction that each cell-type contributes to the total in each sample. (B) For each methylation locus (rows), the methylation levels in METSIM samples or for different cell-types (columns) are shown in the heatmap, the color represents the methylation levels. (C) Correlation between cell-type composition and clinical trait. Traits are plotted in rows, and cell-types are plotted in columns. The color in the heatmap represents spearman correlation between the fraction derived from each cell-type and a clinical trait for and an individual, across all individuals in the METSIM samples.

Since highly expressed genes are often correlated with lower methylation levels in their promoters, we hypothesize that if our genes with significant associations with metabolic syndrome are expressed in adipose tissue, they will also have lower methylation levels in the cell types in which they are expressed. When we examine DNA methylation levels in CpG regions associated with traits in our EWAS, we find that adipocytes tend to have lower methylation levels at these loci, relative to other cell types, suggesting that many of them may be specific to adipocytes. However, two associated loci at *RXRA* and *RAP1GAP2* genes also show decreased methylation levels in macrophages and neutrophils, suggesting that DNA methylation at these loci may be derived from macrophages and/or neutrophils.

Finally, although the relative content of cell-types such as macrophages may be small, they can still contribute to expression and methylation levels, and to clinical phenotypes. We studied the correlation between the cell-type content and clinical traits across all individuals, and found that macrophage content was positively correlated with the clinical traits associated in our EWAS (Figure 4C). These results support the notion that both adipocytes and macrophages contribute to DNA methylation signatures, and to associations between DNA methylation and clinical traits. The correlation between neutrophil content and traits is minimal, suggesting that methylation levels derived from neutrophils are potentially derived from blood contamination during collection of biopsies.

### CHORMATIN STATES AT CANDIDATE LOCI

We used the Roadmap (37) and RegulomeDB databases to examine chromatin marks and chromatin states in adipocytes or adipose tissue in each of the EWAS loci. The chromatin marks found in each locus are summarized in Figure S4. The sites group into three clusters. The first represents regions of active transcription that contain H3K36me3. The third cluster likely contains enhancers, which are marked by H4K4me1 and H3K27ac. The second cluster is more heterogeneous and has generally fewer marks, with a few sites showing no marks at all.

### DNA METHYLATION BIOMARKER FOR TYPE II DIABETES

DNA methylation is a useful biomarker for assessing the age (38, 39) and BMI (4) of an individual. We asked whether we could develop a biomarker for adipose tissue that could be used to assess a metabolic health outcome, type II diabetes. To this end, we first developed an aggregate measure of type II diabetes by combining multiple clinical traits measured in the METSIM cohort using principal component analysis. Briefly, we split phenotype data into training (n = 6,103) and testing (n = 4,069) sets. We selected traits for inclusion into the aggregate measure of metabolic health using a greedy algorithm that considered combinations of features that produced the largest Welch’s test statistic in the first principal component, when comparing healthy individuals to individuals who had received a type II diabetes diagnosis at baseline examination in the training dataset. Our final measure consists of a linear combination of six traits: two measurements of glucose at baseline and at two hours during an oral glucose tolerance test, a binary measure of elevated blood glucose, two measurements of urine albumin levels at the start and end of collection, and one measurement of LDL levels (Table 2). We decomposed the testing data using the trained linear combination of the six selected features.

**Table 2:** PC1 Feature Contribution

| **Table 2. PC1 Feature Contribution** |  |
| --- | --- |
| Variance Explained = 0.295 |  |
| **Feature** | **Contribution** |
| Glucose Baseline | 0.660 |
| Glucose 120min | 0.054 |
| Elevated Blood Glucose | 0.191 |
| LDL | -0.466 |
| Urine Albumin Baseline | 0.402 |
| Urine Albumin 60min | -0.367 |

This first principal component allows us to effectively segregate individuals by type II diabetes status as baseline (Figure 5A). A follow-up examination was conducted on METSIM participants an average of 53.2 months (std = 12.6) after the baseline examination. This allows us to identify a subpopulation that was healthy at baseline but develops type II diabetes at follow-up. Based on the PC1 score this group has baseline levels that are intermediate between the healthy and type 2 diabetes group (Figure 5A). This suggests that our approach is also able to detect individuals at risk of developing type II diabetes. Additionally, the first principal component outperforms individual metabolic metrics commonly used for diagnosis of type II diabetes (40) in the classification of type II diabetes status at baseline or follow-up examination (Figure 5B). These results suggest that PC1 is a useful metric for assessing risk of developing type II diabetes.

**Figure 5:**
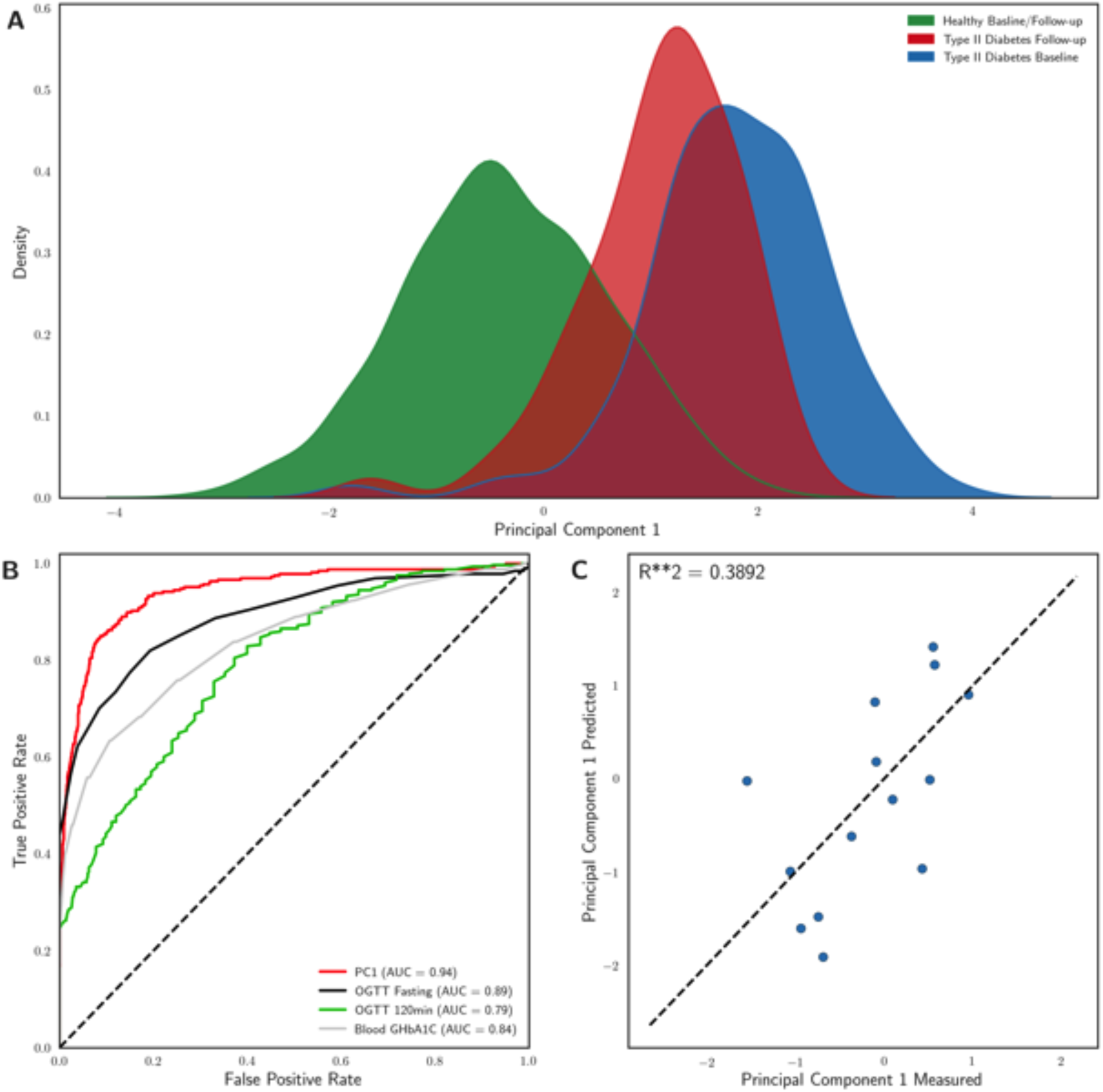
A methylation biomarker to assess type II diabetes. (A) Kernel density estimates of principal component 1 by type II diabetes status at baseline and follow-up examination for METSIM participants who received follow-up examination in the testing dataset (n = 2422). Healthy individuals had not been diagnosed with type II diabetes at baseline or follow-up examination, type II diabetes follow-up individuals had not received a type II diabetes diagnosis at baseline but were diagnosed by follow-up examination, and type II diabetes baseline individuals received a type II diabetes diagnosis before or at baseline examination. (B) A combined type II diabetes feature, PCI, outperforms individual features for classification of diabetes at baseline or follow-up examination among METSIM participants in the testing dataset who received follow-up examination (n = 2422). (C) Measured and predicted PC1 values for a single cross-validated regression model fit to 18 CpG sites.

Finally, we asked whether we could predict the value of PC1 using DNA methylation, in order to develop a biomarker to assess type II diabetes risk. We split methylation data into a training set (n = 213) and a testing set (n = 15) ran the model separately three times. We used randomized lasso to select CpG sites used to generate a linear model that predicts the PC1 value of each individual, and selected 24 CpG sites across the runs (supplemental file). We used a five-fold cross validation approach to fit a model with the training data. We measured the accuracy of this approach using the testing data across three separate runs, and found that the average R-squared between our predicted and measured PC1 values was 0.4034. This suggests that using a subset of CpG sites measured in adipose tissue we are able to predict the risk of developing diabetes.

## DISCUSSION

In this study, we utilized natural variation in DNA methylation in the adipose tissue of a human population to explore the relationship between DNA methylation and complex clinical traits associated with metabolic syndrome. We chose to focus our analysis on adipose tissue, as it is believed to be the central tissue mediating metabolic syndrome traits. While metabolic syndrome surely involves a complex interplay between adipose tissue, liver and immune cells, it is likely that adipose tissue undergoes the most dramatic epigenetic changes during the advancement of metabolic syndrome. In fact, previous studies have shown that adipose tissue has significant epigenetic differences between lean and obese individuals (41).

Using epigenome-wide analysis we identified 21 novel associations for diabetes and obesity phenotypes, corresponding to 24 candidate genes. We further narrowed our candidates to 18 high confidence candidate genes based on presence of *cis*-eQTL for these genes in adipose tissue (Table 1). Our results demonstrate the power of EWAS to identify significant associations for metabolic traits in humans using only 201 individuals, and highlight how epigenetic factors such as DNA methylation could be considered in conjunction with genetic variation to elucidate the complex cellular mechanisms that ultimately lead to observable phenotypes.

We found three loci where multiple clinical traits mapped to the same methylation region and associated gene, *SLC1A4* on chromosome 2 (Figure 3), *CPEB4* on chromosome 5 (Table 1), and *FASN* on chromosome 17 (Figure 2). *FASN* is a known regulator of fatty acid metabolism (42), and its expression is associated with mature adipocytes replete with the downstream products of FASN. We observed that DNA methylation in the *FASN* gene was correlated with multiple metabolic syndrome clinical traits. Remarkably, we show that the variation in DNA methylation levels of FASN capture approximately 16% of the variation of metabolic traits such as BMI (Figure 2D), a significant portion of the variation in the trait in our population.

The mechanisms by which metabolic syndrome traits affect methylation levels are still incompletely understood. While transcriptional levels of *FASN* and other genes likely respond quickly to insulin release, it is well established that, in contrast, DNA methylation levels are very stable, and change on a much slower timescale. We expect that methylation levels in the body of a gene change on the timescale of weeks, in response to the daily changes of insulin signaling. Thus, we hypothesize that DNA methylation levels at or near genes may reflect the history of insulin signaling during the previous weeks, and are thus robust markers for the average physiological state of the individuals. We observed that the expression of *FASN* decreases with increasing obesity, in agreement with previous multiple studies showing that obese individuals have lower insulin sensitivity (43), leading to lower *FASN* expression. The *FASN* regions we identified in our study are intragenic, and associated with several histone marks including H3k36me3, and H3k4me3 and H3k4me1. These marks are often associated with the boundaries of promoters and transcribed regions. Thus, we speculate that the reduced insulin sensitivity leads to a hypermethylation of the promoter, which is associated with a decrease in gene expression.

We also observed that the methylation levels near *RXRA* are associated with metabolic traits. RXRA is known to form a complex with PPARG, a master regulator of adipogenesis and adipocytes (44). In contrast to *FASN* and *RXRA*, the amino acid transporter *SLC1A4*, previously linked to metabolite levels and atherosclerosis (45), is a novel gene associated with both diabetes and obesity traits. The cytoplasmic polyadenylation element binding protein 4, *CPEB4* has been previously associated with obesity (46), and waist-to-hip ratio (47), but not with diabetes. Here we find that *CPEB4* is associated with measures of insulin sensitivity/ insulin resistance.

We found several other novel gene associations such as *LINC01317* which we found to be associated with insulin sensitivity (MATSUDA index) and *TPCN1* which was associated with body weight in our EWAS. In addition, we found Strawberry Notch Homolog 2 *(SBNO2)* to be associated with BMI and body weight in our study, and with BMI in a previous EWAS (4). *SBNO2* regulates inflammatory responses (48), and a *Sbno2* mutant mouse model shows impaired osteoclast fusion, osteoblastogenesis, osteopetrosis, and increased bone mass (49).

As adipose tissue is a heterogeneous tissue that contains adipocytes, endothelial and immune cells, among others, we asked whether we could determine which cell types were most significant in our analyses. We found that most of our significantly associated loci were hypomethylated in adipocytes compared to other cell types. This suggested that most of the methylation variation we observe is likely occurring in adipocytes, which constitute the majority of cells in adipose tissue. Using a DNA methylation based deconvolution approach we also estimated the abundance of each cell type in each individual. As expected, we found that adipocytes constituted around 80% of the cells in our samples. Intriguingly, however, we observed that the abundance of macrophages varied across individuals in a manner that was strongly correlated with metabolic traits. This suggests that obese individuals have higher macrophage counts in their adipose tissue compared to lean individuals, a result that supports pervious observations (36).

In previous studies DNA methylation has been used to develop robust biomarkers for multiple traits such as age (38) and BMI (4). We therefore asked whether we could develop an accurate biomarker to assess type II diabetes risk from our data. We first aggregated clinical traits to define a metric of metabolic health that is associated with the risk of developing type II diabetes, by combining measures of glucose, LDL and urine albumin. We showed that this metric stratifies the population into healthy and diabetic individuals. We also showed that high values of this metric strongly associated with the development of type 2 diabetes in follow-up measurements in the METSIM cohort. Finally, using a limited set of CpG sites, we developed a model that accurately predicts the values of this metric. This result suggests that DNA methylation measurements in adipose tissue can be used to assess the risk of developing type II diabetes. It is important to note that an adipose tissue biomarker may not be practical for clinical use as it necessitates the use of adipose tissue biopsies. Additional work is required to verify whether the biomarker translates to clinically relevant tissue types such as blood.

The current study outlines the usefulness of examining epigenetics in a disease relevant tissue, but is constrained by limited genetic variability of the METSIM cohort. The METSIM cohort is composed of middle-aged Finnish men, and it is likely that some of our results will not extend to other ethnic populations or to female cohorts. Global methylation patterns are known to differ between males and females in blood (50), and these sex specific methylation differences and their relationship with metabolic traits should be explored in future work. Furthermore, an open question is how the epigenetic profile varies between multiple tissues under the same physiological conditions. Comparing DNA methylation data from tissues such as liver, muscle, and visceral adipose may elucidate how these respond differently to metabolic syndrome. Finally, our study population was enriched for healthy individuals, future work should focus on the epigenetic differences between diabetic/insulin insensitive individuals and healthy individuals.

In conclusion, our DNA methylation profiles of adipose tissue allowed us to identify loci that are likely reacting to the metabolic state of an individual, but whose modulation is also likely to affect the individual’s metabolic profile. Our results demonstrate the usefulness of utilizing population variation in DNA methylation for identifying genes associated with complex clinical traits. Here, we identified 18 novel candidate genes for metabolic syndrome using the adipose tissue of 201 individuals. None of these loci could be found using GWAS in 152 individuals of the same cohort. Since DNA methylation in a fraction of CpGs is heritable and regulated by genetics in *cis* and in *trans* (3, 17, 51), EWAS and GWAS can be used in a complementary manner to uncover heritable factors contributing to the etiology of complex traits.

## MATERIALS AND METHODS

### Data access

RRBS sequencing data and all EWAS association results can be obtained from GEO: GSE87893.

### Clinical phenotypes on human subjects

Ethics Committee of the Northern Savo Hospital District approved the study. All participants gave written informed consent. Clinical trait phenotypes for the EWAS study were collected on 201 individuals from the METSIM cohort (19, 20, 35). The population-based METSIM study included 10,197 men, aged 45 to 73 years, from Kuopio, Finland. After 12 hours of fasting, a 2 hour oral 75 g glucose tolerance test was performed and the blood samples were drawn at 0, and 120 min. Plasma glucose was measured by enzymatic hexokinase photometric assay (Konelab Systems reagents; Thermo Fischer Scientific, Vantaa, Finland), and insulin and pro-insulin were determined by immunoassay (ADVIA Centaur Insulin IRI no. 02230141; Siemens Medical Solutions Diagnostics, Tarrytown, NY, USA). Plasma levels of lipids were determined using enzymatic colorimetric methods (Konelab System reagents, Thermo Fisher Scientific, Vantaa, Finland). Plasma adiponectin was measured with Human Adiponectin Elisa Kit (Linco Research, St Charles, USA), C-reactive protein (CRP) with high sensitive assay (Roche Diagnostics GmbH, Mannheim, Germany), and interleukin 1 receptor agonist (IL1RA) with immunoassay (ELISA, Quantikine DRA00 Human IL-1RA, R&D Systems Inc., Minneapolis, USA). Serum creatinine was measured by the Jaffe kinetic method (Konelab System reagents, Thermo Fisher Scientific, Vantaa, Finland) and was used to calculate the glomerular filtration rate (GFR). Height and weight were measured to the nearest 0.5 cm and 0.1 kg, respectively. Waist circumference (at the midpoint between the lateral iliac crest and lowest rib) and hip circumference (at the level of the trochanter major) were measured to the nearest 0.5 cm. Body composition was determined by bioelectrical impedance (RJL Systems) in participants in the supine position. Summary statistics for each phenotype are shown in Table S1. We then transformed the residuals using rank-based inverse-normal transformation for downstream analyses. This transformation involves ranking a given phenotype’s values, transforming these ranks into quantiles and, converting the resulting quantiles into normal deviates. The goal of this transformation is to minimize spurious associations due deviations from the underlying assumption that data are normally distributed, it is common practice for GWAS of quantitative traits (32).

### Evaluation of insulin sensitivity

We evaluated insulin sensitivity by the Matsuda index and insulin resistance by the HOMA-IR as previously described (20).

### RRBS libraries

We prepared genomic DNA from adipose tissue biopsies with the DNeasy extraction kit (Qiagen, Valencia, CA, USA). We prepared RRBS libraries as previously described (17, 52), with minor modifications. Briefly, we isolated genomic DNA from flash frozen adipose biopsies using a phenolchloroform extraction, digested 500ng of DNA with MspI restriction enzyme (NEB, Ipswich, MA, USA), carried out end-repair/adenylation (NEB) and ligation with TruSeq barcoded adapters (Illumina, San Diego, CA, USA). We selected DNA fragments of size range 200-300bp with AMPure magnetic beads (Beckman Coulter, Brea, CA, USA), followed by bisulfite treatment on the DNA (Millipore, Billerica, MA, USA), and PCR amplification (Bioline, Taunton, MA, USA). We sequenced the libraries by multiplexing 4 libraries per lane on the Illumina HiSeq2500 sequencer, with 100bp reads.

### Sequence alignment

We aligned the reads with BSMAP to the hg19 human reference genome (27). We trimmed adapters with Trim Galore! (www.bioinformatics.babraham.ac.uk/projects/trim_galore/), allowed for up to 4 mismatches and selected uniquely aligned reads. We have previously shown that BSMAP performance is comparable to the BS-Seq aligners BS-Seeker2, and Bsmark in terms of accuracy and mappability (53).

### Methylation regions

We filtered aligned data to keep only CpG cytosines with 10x coverage or more across all samples, and with data coverage in at least 75% of the samples. This resulted in 2,320,297 CpGs. From these cytosines, we defined methylation regions grouping nearby cytosines together in expanding windows using the following rules: 1) We treated each cytosine as a seed cytosine for a potential methylation region and in order to expand the region methylation levels were required to be correlated across the cohort at an R2> = 0.9 for directly adjacent cytosines, 2) extension of the methylation region from the seed cytosine was allowed to continue up to 500bp in either direction from the seed cytosine, 3) after region extension, only regions with greater than 2 cytosines were retained, 4) overlapping or adjacent regions remaining from (3) were then merged, 5) a methylation region was limited to 3kb maximum. We chose these parameters since the majority of CpG islands are less than 3kb in size (54). Methylation for resulting regions was calculated as the average methylation of all included cytosines. The distribution of region size we observed was a minimum of 3bp, maximum of 3kb, and median 143bp.

### EWAS

We used the linear mixed model package pyLMM (https://github.com/nickFurlotte/pylmm) to test for association and to account for potential population structure and relatedness among individuals. This method was previously described as EMMA (33), and we implemented the model in python to allow for continuous predictors, such as CpG methylation levels that vary between 0 and 1, as previously described (17). We applied the model: y = μ+xβ+u+e, where μ=mean, x=CpG, β=CpG effect, and u=random effects due to relatedness, with Var(u)=σ_g_^2^K and Var(u)=σ_e_^2^K, where K=IBS (identity-by-state) matrix across all CpG methylation regions. We computed a restricted maximum likelihood estimate for σ_g_^2^K and σ_e_^2^K, and we performed association based on the estimated variance component with an F-test to test that β does not equal 0. Associations were considered significant if the *p*-value for the association was below 1×10^-7^, based on the Bonferroni correction for the number of CpG regions tested.

### Inflation

We calculated the inflation factor lambda by taking the chi-squared inverse cumulative distribution function for the median of the association *p*-values, with one degree of freedom, and divided this by the chi-squared probability distribution function of 0.5 (the median expected *p*-value by chance) with one degree of freedom. We plotted qqplots for representative phenotypes using the *qqplot* function in Matlab, with a theoretical uniform distribution with parameters 0,1.

### Adipose expression from human subjects

Expression levels from adipose tissue biopsies were collected on 770 individuals of the METSIM cohort as previously described (19, 35), and 151 of these subjects were also represented in the current methylation dataset. Total RNA from METSIM participants was isolated from adipose tissue using the Qiagen miRNeasy kit, according to the manufacturer’s instructions. RNA integrity number (RIN) values were assessed with the Agilent Bioanalyzer 2100 instrument and 770 samples with RIN >7.0 were used for transcriptional profiling. Expression profiling using Affymetrix U219 microarray was performed at the Department of Applied Genomics at Bristol-Myers Squibb according to manufacturer’s protocols. The probe sequences were re-annotated to remove probes that mapped to multiple locations, contained variants with MAF > 0.01 in the 1,000 Genomes Project European samples, or did not map to known transcripts based on the RefSeq (version 59) and Ensembl (version 72) databases; 6,199 probesets were removed in this filtering step. For subsequent analyses, we used 43,145 probesets that represent 18,155 unique genes. The microarray image data were processed using the Affymetrix GCOS algorithm using the robust multiarray (RMA) method to determine the specific hybridizing signal for each gene.

### PEER factor analysis

We corrected RMA-normalized expression levels for each gene using probabilistic estimation of expression residuals (PEER) factors (55). PEER factor correction is designed to detect the maximum number of *cis*-eQTL. We then transformed the residuals using rank-based inverse-normal transformation. We used the inverse normal-transformed PEER-processed residuals after accounting for 35 factors for downstream eQTL mapping.

### *cis*-eQTL in adipose expression

eQTL studies from the adipose biopsies of the METSIM cohort have been previously described (19). Briefly, gene expression in 770 adipose biopsy samples from the METSIM cohort was measured with Affymetrix U219 microarray. SNP genotyping was performed with Illumina OmniExpress genotyping chip and imputed based on the Haplotype Reference Consortium reference panel. Association of gene expression and SNPs were calculated with FaST-LMM. eQTL were defined as *cis* if the peak association had a *p*-value of *p* < 2.46 × 10^-4^ corresponding to 1% FDR, and if it was found within 1 Mb on either side of the exon boundaries of the gene, as previously described (32).

### GWAS for methylation loci

We performed GWAS on the same 32 clinical traits, transformed using inverse normal transformation as described above, and 681,803 genotyped SNPs for 152 METSIM individuals where we had both genotypes and methylation data. We used a linear model and the R package MatrixEQTL to perform the association, and selected associations where the *p*-value was below 1×10^-7^.

### Published histone marks

We used the Roadmap ChIP-seq datasets to look for any histone marks in human adipocyte and adipose tissue samples. We used the RegulomeDB database to look for evidence of transcription factor footprinting, positional weight matrices (PWM), and active transcription. We accessed the public datasets at (http://www.roadmapepigenomics.org/data/) and (http://www.regulomedb.org/).

### Published GWAS hits

Literature evidence for GWAS hits in candidate genes was obtained from the NHGRI GWA Catalog.

### DNA methylation deconvolution

To estimate the methylation contribution of different leukocytes to the adipose tissue, we used cell-specific methylation markers from DNA methylation signatures across different cell types. Cell-specific CpG methylation loci were identified from purified leukocyte (macrophages, neutrophils, B cells, CD4 + T cells, CD8 + T cells, NK cells) methylation profiles from the Blueprint epigenome project (56). Since there was only one purified adipocyte primary cell line reference available, we also included the average methylation profile across all adipose samples using in this study, which ostensibly consists primarily of adipocytes. The purified adipocyte cell reference was used as a filter to select cell type-specific CpG loci that are hypomethylated in both the adipocyte cell reference, and in the mean of the methylation levels for the 201 adipose samples. We filtered all cell methylomes to CpG loci that are common between the reference methylation profiles and the METSIM adipose tissues samples. To determine cell-specific methylation across all references, we first used a sliding window to aggregate the methylation profiles into regions of CpG loci with similar methylation (within 40% methylation difference across neighboring CpG within 500bp). Regions were selected that were uniquely hypomethylated for each cell types to provide 279 cell-specific hypomethylated regions. To estimate the proportion of each cell type within samples, we performed a non-negative least squares regression (57) on methylation at the cell-specific regions.

### Aggregate measure of metabolic health

The METSIM cohort metabolic phenotype data included 10,197 individuals, and 484 traits. We dropped individuals with a type 1 diabetes (n=25) diagnosis from further analysis, leaving 10,172 individuals. We processed numeric data for downstream analysis by dropping traits with greater than 10% of data points missing. We imputed missing values using a k nearest neighbors (kNN) approach. KNN imputation of phenotype data occurred as follows: (1) neighbors were ranked on Euclidean distance, and (2) missing values were assigned the average value of the nearest neighbor (k=5). Following imputation, we scored phenotype data for normality (scipy.stats.normaltest) (58, 59). We designated the threshold for a normally distributed trait by randomly simulating normally distributed data of equal length as the METSIM phenotype data 1000 times, scoring the random distribution, and setting the threshold at the 90th percentile score of the simulated distributions. We normalized traits following a normal distribution (mean=0, STD=1), and used rank based inverse normalization (mean=0, STD=1) for traits from a non-normal distribution.

We manually removed traits directly predictive of type 2 diabetes status, such as family history or metformin consumption. Following data normalization, we held out 40% of the samples (n=4,069) with phenotype information from feature selection, including samples with RRBS, for downstream analysis. We screened features for the remaining 60% of the samples (n=6,103) for incorporation into the metatrait using randomized logistic regression model (60). We evaluated selected features on their ability to distinguish between individuals with and without type 2 diabetes at baseline using Welch’s t-test. Starting with the feature that had the highest Welch’s test statistic, we considered combinations of traits by iterating through all traits that passed the initial screen, incorporating the trait with the starting trait, performing PCA on the combined traits, scoring the 1st principal component using Welch’s t-test, and returning the set of traits that produced the largest Welch’s test statistic. We repeated the process until Welch’s test statistic no longer increased. We selected six features for incorporation into the meta-trait (Table 2). We decomposed the trait matrix for the test samples using the trained linear combination of selected features.

We implemented trait analysis pipelines in Python3.6.1, utilizing scikit-learn-0.18.1 (61), numpy-13.1 (62), scipy-0.19.1 (63), pandas-0.20.3(64), seaborn-0.8.1(65), and matplotlib-2.0.1 (66) packages.

### Metabolic syndrome biomarker model

We pre-processed the methylation matrix for model fitting by dropping all CpG sites with greater than 10% of data points missing. We imputed missing values using a kNN sliding window approach. We replaced individual CpG sites with missing data with the average value of the 5 nearest neighbors by Euclidean distance within a 6Mb window. The resulting matrix contained 1,633,360 CpG sites. To speed up processing time we only considered CpG sites with variation greater than 0.05. We then split the complete methylation matrix into a training set and a testing set by randomly selecting training samples across the PC1 distribution. Samples were placed into 6 equally sized bins, 95% of samples in each bin were selected for training, resulting in 215 training samples and 17 testing samples. Using the training dataset, we selected CpG sites using randomized lasso regression implemented in scikit-learn-0.19.1. To control for cell type composition differences between samples when selecting CpG sites we decomposed he methylation matrix using PCA, and then reconstructed the methylation matrix without the top three principal components, a method previously shown to control for cell type composition.(67) We performed randomized lasso regression for multiple subsets (n=100) composed of 90% of the training samples and selected CpG sites selected in greater than 40% of the runs. Twenty-four CpG sites were selected across three separate runs. We utilized a five-fold cross validation strategy for model fitting on the selected CpG sites. The fit model was then used to calculate predicted PC1 values for the held out testing samples. The final model consists of the average coefficient for each CpG sites and the average intercept across all cross-validated models. We annotated CpG sites with GREAT-3.0.0 (68) to generate a list of index genes. See supplemental file 1 for a list of CpG sites, regression coefficients for the biomarker model, and index genes.

### Code Repository

Custom code used in data processing and analysis can be found at https://github.com/NuttyLogic/METSIM_HMG_Code.

## ACKNOWLEDGEMENTS

L.D.O. was supported by the Ruth L. Kirschstein National Research Service Award T32AR059033.

M.P and A.J.L were supported by National Institutes of Health (NIH) grant HL28481.

A. J.L. was supported by NIH grants HL30568 and 1P50 GM115318.

M.L. was supported by grants from Academy of Finland and Juselius Foundation.

K.L.M. was supported by NIH grant R01DK093757.

S.E.J. is an Investigator at the Howard Hughes Medical Institute.

## Supplemental Text

### OVERLAP OF EWAS RESULTS AND PUBLISHED GWAS AND EWAS

To further examine the role of candidate genes in metabolic syndrome, we queried published GWAS results in the GWAS catalog (1) to determine if any of our candidate genes had been previously bookmarked a candidate gene in a GWAS for cardiovascular or metabolic traits. Overall, we found 7 candidate genes associated with EWAS hits in the current study have been previously associated with metabolic or cardiovascular. For example, we found that *LINC01317* is associated with insulin sensitivity index (MATSUDA) using EWAS, and a GWAS to Glomerular filtration rate reported a significant association with a SNP (rs10495809) in an intron of *LINC01317* (2). *CPEB4* is associated OGTT plasma insulin levels, insulin sensitivity index (MATSUDA), and insulin resistance index (HOMAIR) in the current EWAS, and was previously reported as a bookmark gene for a SNP ((s7705502) reported in a GWAS to waist-to-hip ratio adjusted for body mass index (3). *CPEB4* is also a strong c/s-eQTL in adipose tissue (Table1, *p*=2.4×10^-174^), making it an excellent candidate gene associated with diabetes in humans. Although the traits associated with the candidate genes are different in the current EWAS and published GWAS, these findings lend additional support to the involvement of these candidate genes in diabetes traits. Results for all candidate genes are summarized in Table S2.

In addition, to published GWAS associations, four of the candidate genes have been previously shown to influence diabetes and/or obesity, including the fatty acid synthase gene *FASN* and the transcription factor *RXRA*, a well-known regulator of metabolism. The mismatch repair genes *MSH2* and *MSH6* were previously linked to BMI in humans (4), and in the current study they are associated with BMI, waist circumference, and percent fat mass.

A recent EWAS study examined association of BMI and DNA methylation levels measured from blood using methylation arrays, in a human population of 10,774 individuals (5). They identified 278 in CpG sites associated with BMI at (*p*< 1* × 10^-7^), distributed across 207 loci, and replicated 187 of these. The published EWAS and the current study both examine BMI as a trait, however we used both different cell-types (blood in the published work, and adipose tissue in our study), and different methods of measuring methylation levels (microarrays versus bisulfite sequencing in our study). Hence to be able to compare our EWAS results with this published study, we focused on the overlap of the nearest candidate gene to each locus, and found that *SBNO2* was associated with BMI in the published EWAS by Wahl and colleagues, and was associated with both BMI and body weight. Additionally, the disparity in sample size (201 vs 10,774) suggest the current study is underpowered to detect all overlapping signals.

---

**Table S1:** Clinical trait descriptions and lambda

| Clinical trait | Mean | SD | Minimum | Maximum | EWAS<br>Inflation<br>factor $\lambda$ |
| --- | --- | --- | --- | --- | --- |
| Body mass index | 26.65 | 3.59 | 19.49 | 44.49 | 0.93 |
| Glycated Hemoglobin A1C | 5.61 | 0.31 | 4.60 | 6.60 | 1.01 |
| HOMAIR insulin resistance index based on HOMA | 1.95 | 1.28 | 0.44 | 9.53 | 0.90 |
| Plasma adiponectin (ug/ml) | 7.02 | 3.16 | 1.40 | 21.40 | 1.12 |
| Plasma C-reactive protein (mg/l) | 2.14 | 3.75 | 0.14 | 42.54 | 1.24 |
| Plasma IL1 receptor antagonist (pg/ml) | 210.60 | 142.07 | 31.00 | 1460.60 | 0.79 |
| Plasma IL1 beta (pg/ml) | 0.38 | 0.90 | 0.09 | 7.81 | 0.84 |
| OGTT fasting plasma free fatty acids (mmol/l) | 0.36 | 0.14 | 0.11 | 0.89 | 0.76 |
| OGTT fasting plasma glucose (mmol/l) | 5.77 | 0.48 | 4.60 | 7.20 | 0.83 |
| OGTT fasting plasma insulin (mU/l) | 7.48 | 4.58 | 2.00 | 32.00 | 0.97 |
| OGTT 120 min plasma insulin (mU/l) | 47.56 | 51.72 | 3.40 | 280.70 | 0.98 |
| OGTT 30 min plasma insulin (mU/l) | 60.12 | 39.21 | 13.40 | 213.10 | 0.81 |
| OGTT fasting plasma proinsulin (pmol/l) | 13.09 | 6.25 | 4.10 | 44.00 | 0.84 |
| OGTT 120 min plasma proinsulin (pmol/l) | 48.04 | 26.49 | 9.40 | 200.10 | 0.96 |
| Bioimpedance Resistance | 475.23 | 53.96 | 350.00 | 636.00 | 0.90 |
| HDL cholesterol (mmol/l) | 1.51 | 0.42 | 0.86 | 3.75 | 0.80 |
| LDL cholesterol (mmol/l) | 3.46 | 0.73 | 2.12 | 5.44 | 0.82 |
| Total cholesterol (mmol/l) | 5.51 | 0.82 | 3.67 | 8.07 | 0.84 |
| Total triglycerides (mmol/l) | 1.36 | 0.84 | 0.40 | 8.59 | 0.83 |
| Waist/hip ratio | 0.96 | 0.06 | 0.81 | 1.19 | 0.73 |
| Bioimpedance: Reactance | 52.06 | 10.28 | 32.00 | 111.00 | 0.89 |
| Diastolic blood pressure (mmHg) | 88.02 | 8.64 | 64.67 | 118.67 | 0.99 |
| Glomerular filtration rate (eGFR<sub>cockrof</sub>) | 108.78 | 23.51 | 62.74 | 211.92 | 1.08 |
| Fat free mass (%) | 79.89 | 4.04 | 64.60 | 90.40 | 0.99 |
| Fat mass (%) | 20.11 | 4.04 | 9.60 | 35.40 | 1.00 |
| Hip (cm) | 100.50 | 6.46 | 72.00 | 138.00 | 0.87 |
| OGTT insulin area under the curve (pmol/l * min) | 35118.72 | 24130.91 | 7587.00 | 136008.00 | 0.82 |
| MATSUDA insulin sensitivity index (mg/dl, mU/l) | 7.35 | 4.07 | 1.10 | 20.06 | 0.97 |
| Muscle mass (%) | 45.76 | 3.56 | 36.10 | 69.90 | 0.96 |
| Systolic blood pressure (mmHg) | 134.85 | 14.74 | 107.33 | 180.67 | 0.99 |
| Waist circumference (cm) | 96.55 | 10.37 | 74.00 | 146.00 | 0.99 |
| Weight (kg) | 82.76 | 12.48 | 55.00 | 149.00 | 0.99 |

**Table S2:** Overlap between EWAS hits and GWAS for metabolic traits

| EWAS locus | Chr | Bp start | Bp end | Candidate Genes(s) | Gene in GWAS catalog | Phenotype in GWAS catalog | SNP in GWAS catalog | Publication |
| --- | --- | --- | --- | --- | --- | --- | --- | --- |
| **Met1** | 2 | 34545726 | 34545903 | LINC01317, LINC01320 | LINC01317 | Glomerular filtration rate in chronic kidney disease | rs11677592 | (Wuttke *et al.* 2016) |
| **Met2** | 2 | 47979865 | 47980000 | MSH2, MSH6 |  |  |  |  |
| **Met3** | 2 | 65064793 | 65064867 | AFTPH, SLC1A4 | SLC1A4 | Atrial fibrillation | rs2540953 | (Low *et al.* 2017) |
| **Met3** | 2 | 65064793 | 65064867 | AFTPH, SLC1A4 | SLC1A4 | Metabolite levels | rs2160387, rs10211524 | (Kettunen *et al.* 2016), (Shin *et al.* 2014), (Inouye *et al.* 2012) |
| **Met4** | 4 | 7774215 | 7774379 | AFAP1 |  |  |  |  |
| **Met5** | 5 | 173132627 | 173132883 | BOD1, CPEB4 | CPEB4 | Waist-to-hip ratio adjusted for body mass index | rs7705502, rs7705502 | (Shungin *et al.* 2015) |
| **Met6** | 6 | 3132000 | 3132119 | BPHL |  |  |  |  |
| **Met7** | 9 | 137263485 | 137263573 | RXRA | RXRA | Blood pressure | rs4841895 | (Simino *et al.* 2014) |
| **Met8** | 10 | 75600604 | 75600822 | CAMK2G |  |  |  |  |
| **Met9** | 10 | 100183346 | 100183441 | HPS1 | HPS1 | Obesity-related traits | rs6584202 | (Comuzzie *et al.* 2012) |
| **Met10** | 11 | 65248602 | 65248737 | FRMD8, SCYL1 |  |  |  |  |
| **Met11** | 12 | 42883183 | 42883320 | PRICKLE1 | PRICKLE1 | Visceral adipose tissue/ subcutaneous adipose tissue ratio | rs12316797 | (Fox *et al.* 2012) |
| **Met12** | 12 | 113734359 | 113734579 | TPCN1 |  | Response to fenofibrate (adiponectin levels) | rs2384207 | (Aslibekyan *et al.* 2013) |
| **Met13** | 17 | 2893657 | 2893829 | RAP1GAP2 |  |  |  |  |
| **Met14** | 17 | 40913548 | 40913600 | RAMP2 |  |  |  |  |
| **Met15** | 17 | 80041514 | 80042067 | FASN |  |  |  |  |
| **Met16** | 17 | 80046324 | 80047195 | FASN |  |  |  |  |
| **Met17** | 17 | 80050198 | 80050365 | FASN |  |  |  |  |
| **Met18** | 17 | 80051500 | 80053080 | FASN |  |  |  |  |
| **Met19** | 19 | 1148096 | 1148232 | SBNO2 |  |  |  |  |
| **Met20** | 21 | 16661536 | 16661714 | NRIP1, USP25 |  |  |  |  |
| **Met21** | 22 | 28151964 | 28152032 | MN1 | MN1 | Blood pressure smoking interaction | rs133980 | (Sung *et al.* 2015) |
| **Met21** | 22 | 28151964 | 28152032 | MN1 | MN1 | Warfarin maintenance dose | rs68043594 | (Parra *et al.* 2015) |

**Figure S1:**
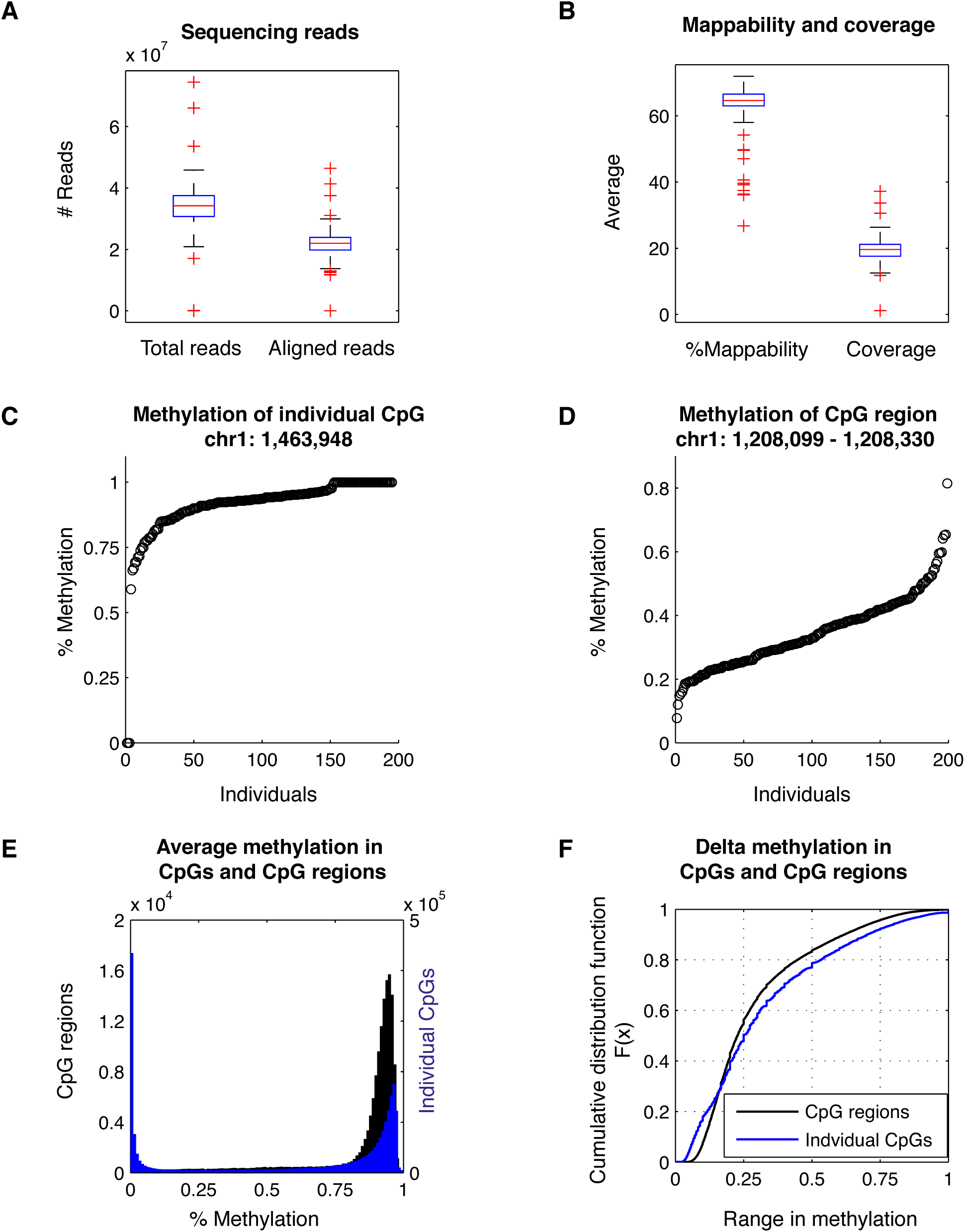
Sample statistics. (A) Total number of reads and aligned reads in all RRBS libraries. (B) Mappability and coverage in all RRBS libraries. (C-D) Methylation patterns for an (C) individual CpG and (D) a CpG methylation region. The x-axes are individual samples, where each dot is a sample. The y-axes are the methylation levels for the CpG in that sample. (E) Histogram showing distribution of average methylation levels for individual CpGs (blue) and CpG regions (black). (F) Distribution of the range in methylation levels across samples for individual cytosines (blue) and methylation regions (black). The x-axis is delta methylation levels across individuals, and the y-axis is the cumulative distribution function F(x).

**Figure S2:**
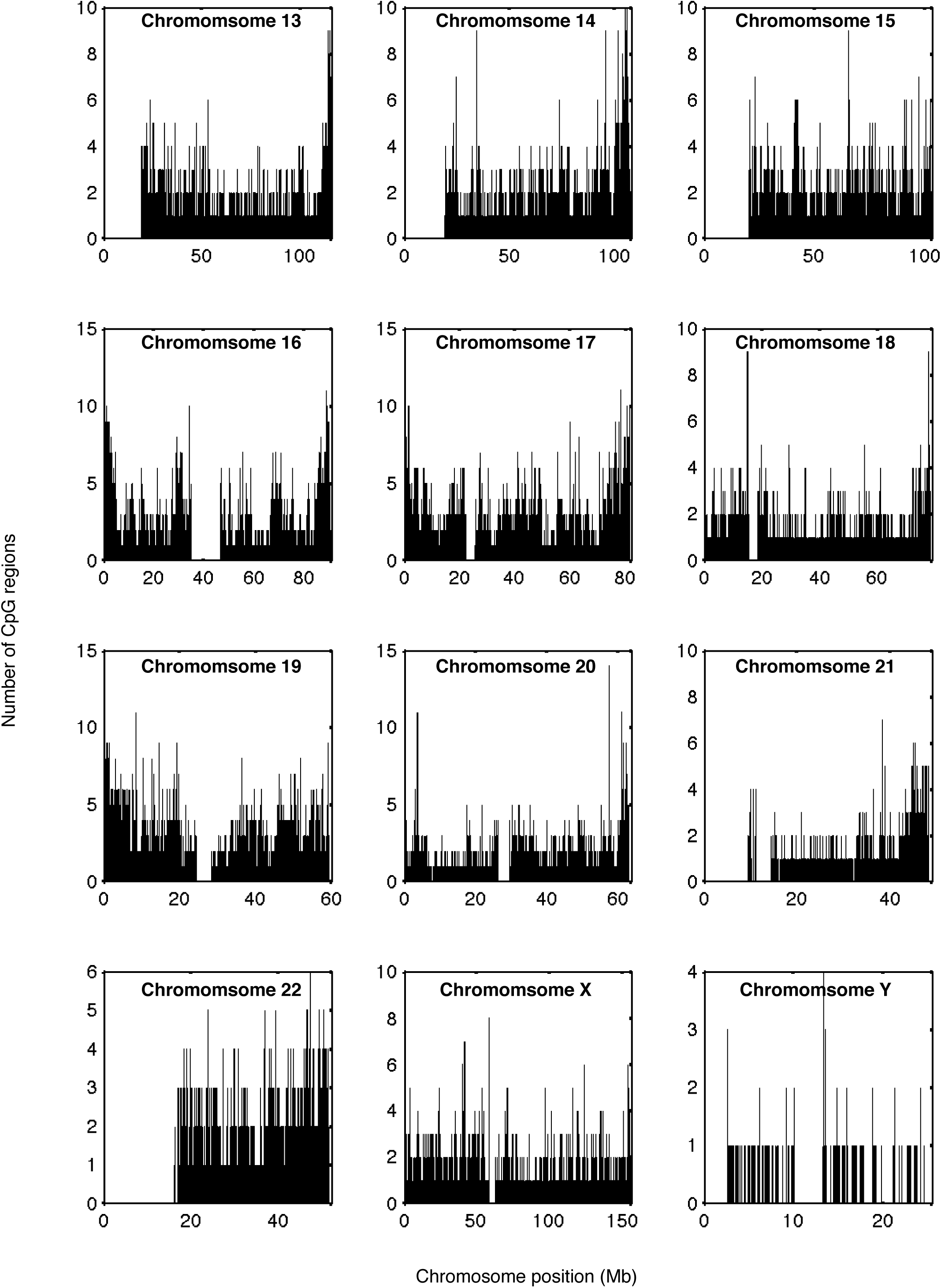
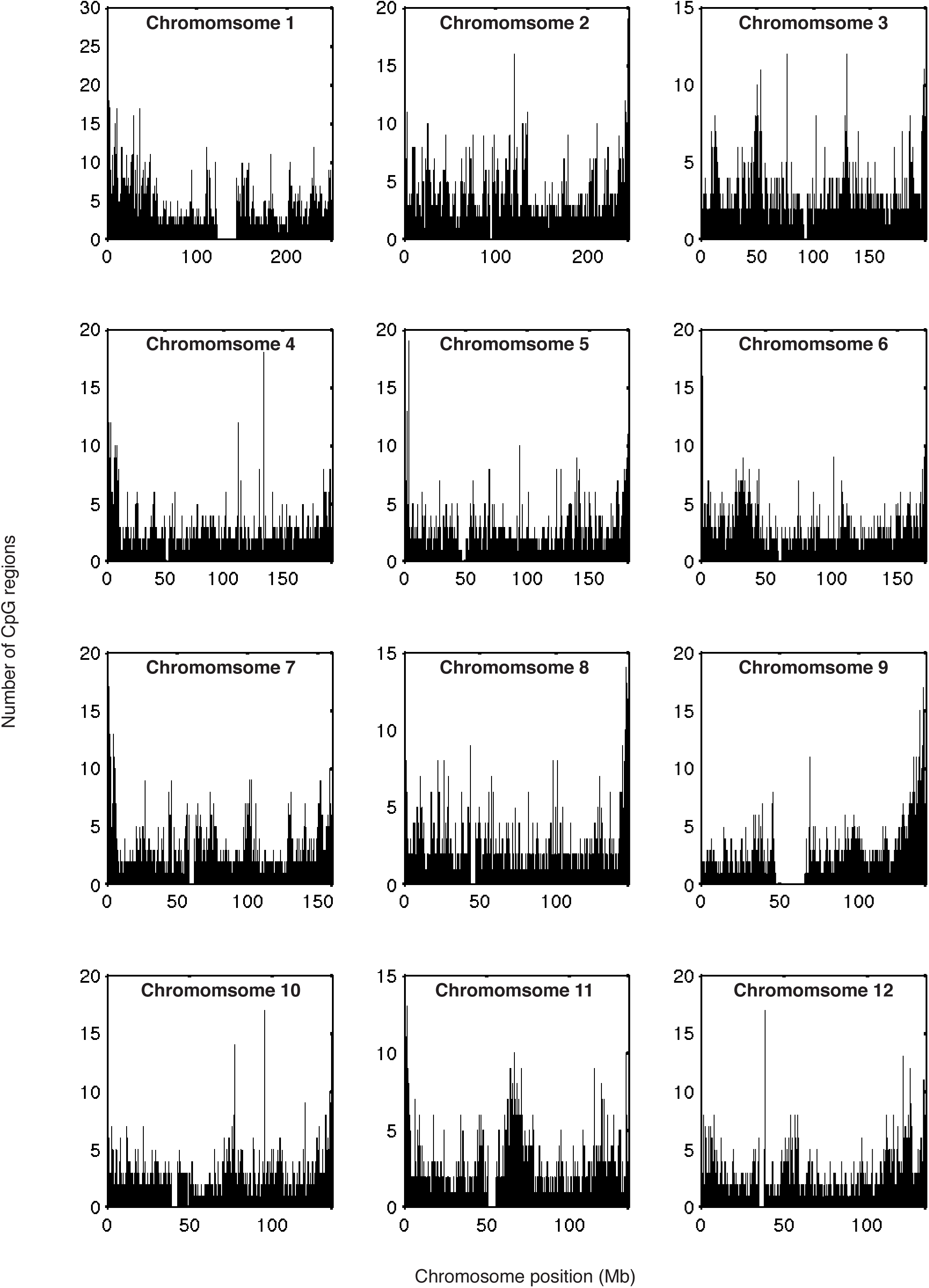
Distribution of CpG regions in human genome.

**Figure S3:**
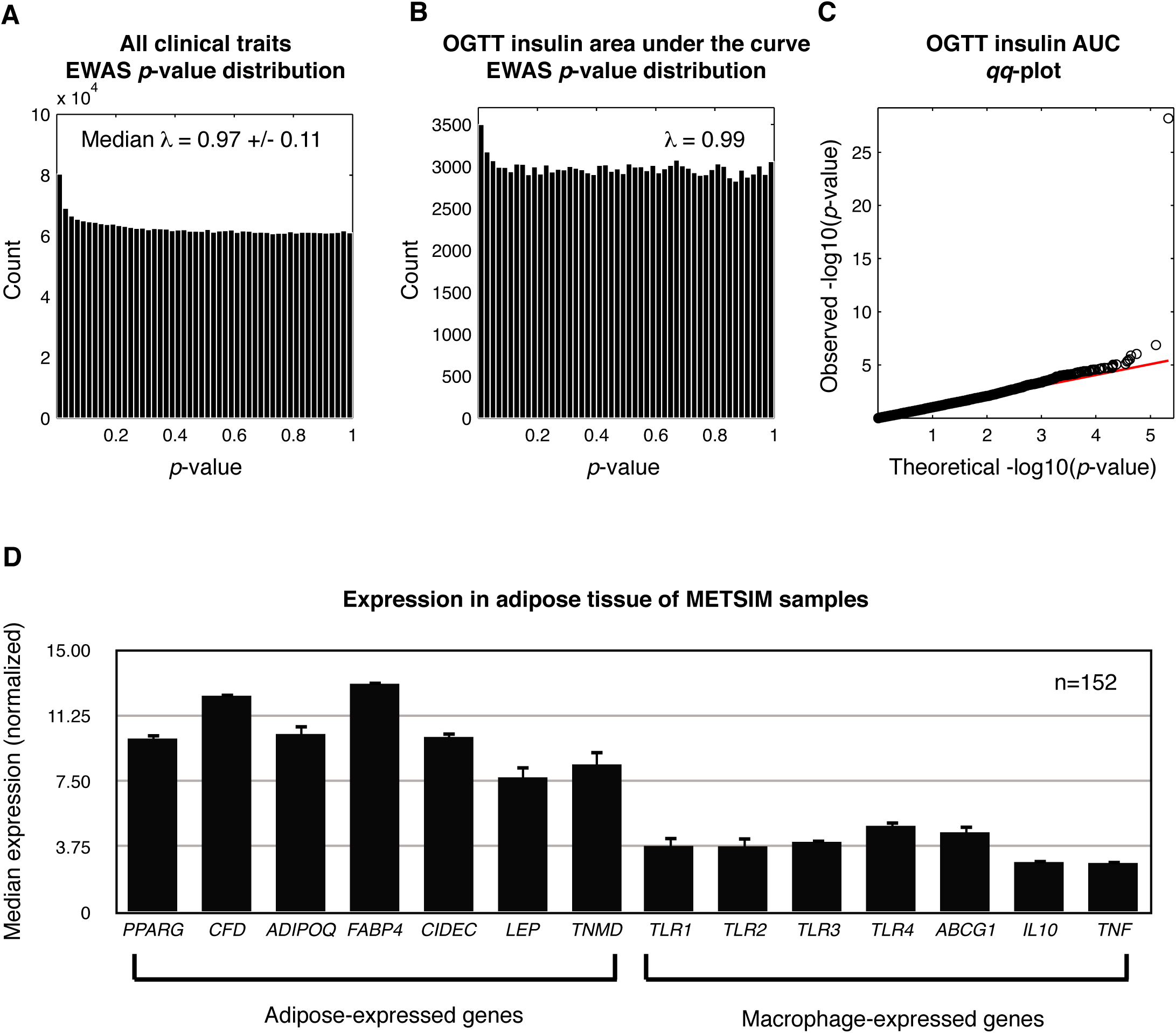
Epigenome-wide assocition *p*-value distribution. EWAS *p*-value distributions and expression of tissue specific genes. Histogram of the *p*-value distributions in the EWAS for (A) all clinical traits, and (B) oral glucose tolerance test area under the curve. (C) Q-Q plot between the theoretical uniform distribution (red) and the EWAS *p*-value distribution (black) for the trait oral glucose tolerance test area under the curve. (D) Expression levels in adipose tissue biopsies for samples where we have both methylation and expression data. The y-axis is the median expression across individuals, and the x-axis are genes known to be expressed in adipocytes or macrophages.

**Figure S4:**
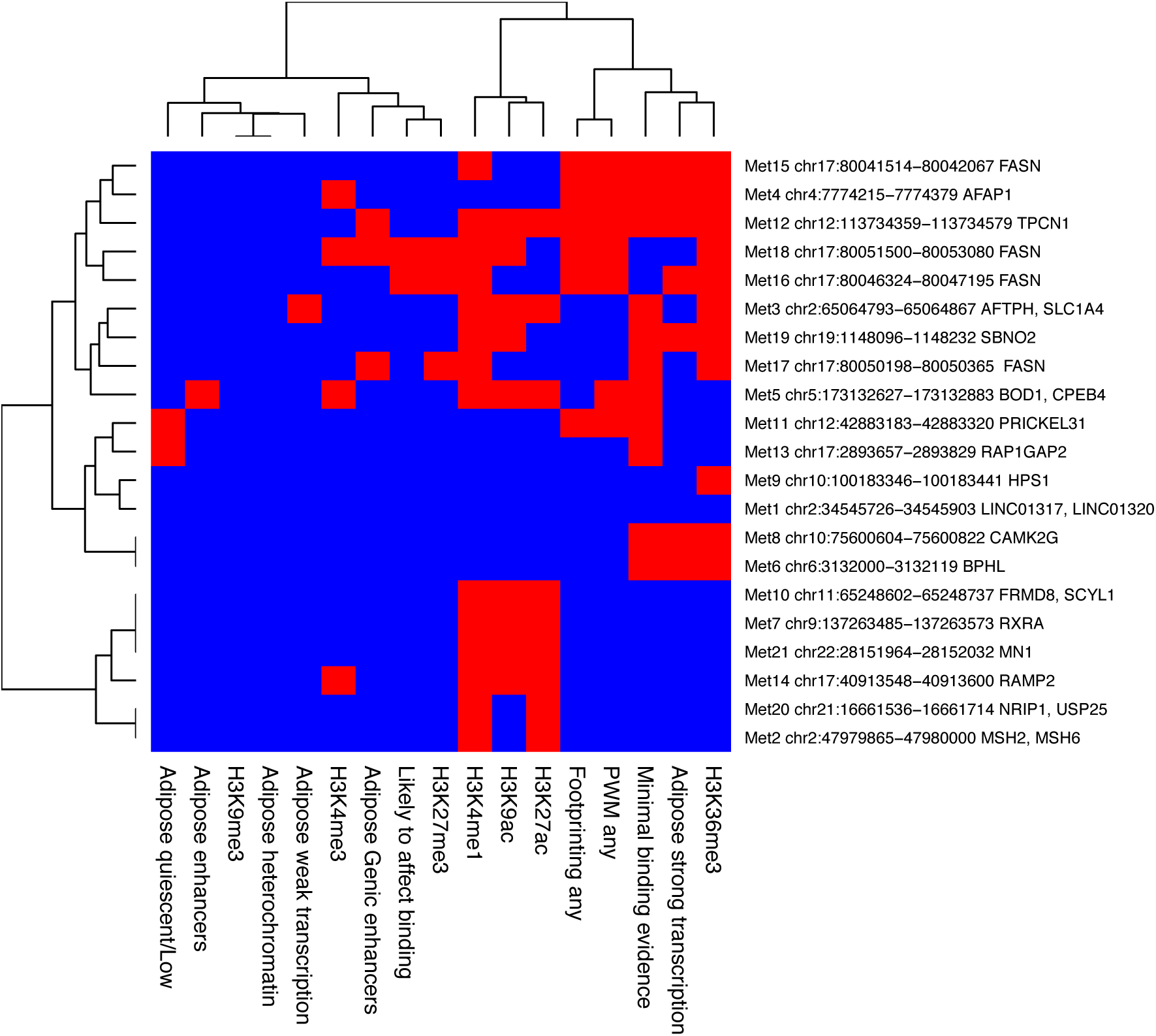
Roadmap and RegulomeDB chromatin marks in loci. Chromatin marks and chromatin states. Chromatin marks and chromatin states for each locus based on Roadmap and RegulomeDB databases. Red=present, blue=absent.

